# Identifying the suite of genes central to swimming in the biocontrol bacteria *Pseudomonas protegens* Pf-5

**DOI:** 10.1101/2023.01.12.523705

**Authors:** BK Fabian, C Foster, A Asher, KA Hassan, IT Paulsen, SG Tetu

## Abstract

Swimming motility is a key bacterial trait, important to success in many niches, including assisting in colonization of host surfaces. Biocontrol bacteria, such as *Pseudomonas protegens* Pf-5 are increasingly being used as an agricultural tool to control crop diseases, where motility is a factor in successful colonization of the plant rhizosphere. Swimming motility has been studied in a range of bacteria and typically involves a suite of flagella and chemotaxis genes, however the specific gene set employed for both regulation and biogenesis can differ substantially between organisms. Here we used transposon directed insertion site sequencing (TraDIS), a genome-wide approach, to identify 249 genes involved in *P. protegens* Pf-5 swimming motility. As expected, flagella and chemotaxis genes comprised a large proportion of these genes. However we also identified a suite of additional genes important for swimming, including genes related to peptidoglycan turnover, O-antigen biosynthesis, cell division, signal transduction, c-di-GMP turnover and phosphate transport, along with 27 conserved hypothetical proteins. Experimental gene knockout mutants and TraDIS data together suggest that defects in the Pst phosphate transporter lead to enhanced swimming motility. Overall, this study expands our knowledge of pseudomonad motility and highlights the utility of a TraDIS-based approach for systematically analyzing the functions of thousands of genes. This work sets a foundation for understanding how swimming motility may be related to the inconsistency in biocontrol bacteria effectiveness and reliability in the field.

**Importance:** Biocontrol bacteria, such as *Pseudomonas protegens* Pf-5 are increasingly being used as an agricultural tool to control crop diseases, and motility is a key factor in their successful colonization of plant surfaces. Here we use a high-throughput approach to identify the suite of genes important for swimming motility in *P. protegens* Pf-5. These included flagella and chemotaxis genes, as well as a variety of cell surface, cell division and signalling genes. We also show that defects in the Pst phosphate transporter lead to enhanced swimming motility, a hitherto unreported link between phosphate transport and swimming motility. Understanding the genetic basis of swimming motility enhances our knowledge of key processes in biocontrol bacteria that are needed to ensure their competitive success. This will contribute to developing strategies to increase the utility of biocontrol bacteria in agricultural settings to prevent crop losses.

## Introduction

The use of biocontrol bacteria as an agricultural tool to control crop diseases is growing, as they can provide many of the same benefits as fertilizers and pesticides without their accompanying damaging effects (1, 2). Colonization of the plant rhizosphere is a crucial step in the provision of the beneficial effects of many biocontrol bacteria (3, 4). Successful colonization requires the movement of bacteria from the site of inoculation to the site of activity, thus motility is an important factor in the field utility of biocontrol bacteria (5-7).

Bacteria use chemotactic systems combined with flagellated movement to detect and move towards more favorable environments (7-9). Swimming is one of the main types of bacterial motility and is characterized by the movement of individual bacteria through an aqueous environment (9, 10) powered by rotating flagella that can be triggered by environmental stimuli (11). Flagella are best studied in the peritrichous bacteria *Escherichia coli* and *Salmonella enterica* ser. Typhimurium, where more than 50 different proteins are involved (12-14), and flagellar synthesis and assembly are regulated via a three-tiered transcriptional cascade (reviewed in Bouteiller et al 2021).

Flagella structural genes are well conserved across Gram negative bacteria, but there are substantial differences in the regulation of flagella synthesis and assembly between the enteric model systems and pseudomonads (15, 16). Pseudomonad species and strains can have differing numbers of flagella, with well-studied strains possessing from one to seven flagella (17, 18). The synthesis and assembly of the *P. aeruginosa* PAO1 single polar flagellum is more complex than production of peritrichous flagella, involving ∼60 genes and regulated by a four-tier hierarchy, comprised of different genes to those of *E. coli* (15, 16).

Outside of well-known flagella and chemotaxis genes, there have been additional genes implicated in bacterial swimming motility (10). Using a *P. aeruginosa* PAO1 Mini-Tn*5*-lux mutant library, seven non-flagellar mutants were identified that had reduced swimming motility, including genes relating to nucleotide metabolism, RNA modification, central intermediary metabolism and four hypothetical genes (19). A transposon insertion sequencing study of *E. coli* EC958 identified 14 non-flagellar genes and intergenic regions involved in enhanced swimming motility, showing that flagellar and chemotaxis genes are only part of the gene suite important for motility (2, 20). Expanding motility research outside of flagella biosynthesis and chemotaxis genes will likely identify additional genes which influence swimming and therefore rhizosphere colonization and biocontrol activity.

The critical role of motility in rhizosphere colonization and biocontrol efficacy has been confirmed in multiple bacterial species (10). In *P. fluorescens* F113, hypermotile mutants are more competitive at rhizosphere colonization than the wildtype strain (4). *P. fluorescens* WCS365 mutants deficient in flagella-driven chemotaxis towards root exudates are poor colonizers of root tips in competition with other bacteria (6). In *P. fluorescens* NBC275, mutants showing a total loss of antifungal activity had reduced swimming motility (21).

Swimming motility is vital for pseudomonads, and biocontrol bacteria more broadly, to be able to colonize the rhizosphere and effectively carry out biocontrol activities.

*Pseudomonas protegens* Pf-5 (hereafter referred to as Pf-5) is a plant growth-promoting bacteria which was originally isolated from the roots of cotton plants (22). Pf-5 produces a range of antibacterial and antifungal secondary metabolites (23) and can control pathogens of a range of crops (24-27). Work on Pf-5 has primarily focused on identifying genes involved in the regulation and production of secondary metabolites (1). Analysis of the Pf-5 genome has revealed many potential genes and molecular systems Pf-5 may utilize in its proliferation in soil and colonization of the rhizosphere (23, 28). Swimming motility studies in Pf-5 have shown that mutations in the global regulators *gacA* and *rpoS* have no effect on swimming, a flagella biosynthesis mutant *flhA* has a deficiency in swimming, and knockouts of the polyurethanases *pueA* and *pueB* have no effect on swimming motility (29-31).

Here we used transposon directed insertion site sequencing (TraDIS), a genome-wide approach, to identify genes involved in Pf-5 swimming motility. TraDIS is a powerful technique that combines high-density random transposon insertion mutagenesis and high-throughput sequencing to study gene fitness and link genotype and phenotype at a genome-wide scale (32, 33). The data from our experiments provides insights into the suite of genes that are important for Pf-5 swimming motility beyond the classic set of swimming related genes.

## Methods

### Bacterial strains and media

*Pseudomonas protegens* Pf-5 was isolated from soil of a cotton field in Texas, USA (22) and a complete genome sequence is available (28; ENA accession number CP000076). Pf-5 was routinely cultured using King’s Medium B at 27°C. Modified KMB agar plates for swimming and the controls contained less proteose peptone (1% w/v), 5.28 mM K_2_HPO_4_, 6 mM MgSO_4_ and used Bacto™ Agar (Becton, Dickinson and Company; 29).

### Transposon mutant library experiments

The role of genes in swimming motility was investigated using a previously constructed *Pseudomonas protegens* Pf-5 transposon mutant library, as described in (35), which contains ∼256,000 unique transposon insertion sites spread evenly throughout the genome, equating to an average of ∼45 transposon insertion sites per non-essential protein coding gene. Two microliters of the Pf-5 transposon mutant library (3.2 × 10^7^ CFUs) were stab-inoculated into the center of six swim agar plates (modified KMB with 0.3% Bacto agar) in triplicate (total of 18 plates). After 24 h at 22°C, 4-5 mm of the agar containing the most motile cells was removed from the edge of the swimming zone (half of the circumference from all six plates; Figure 1). An equivalent volume of KMB was added, and the mixture was vortexed for 10-20 seconds to homogenize the agar. The mixture was pelleted at 16,000 x g for 5 min and excess agar removed (termed the ‘output pool’). For the controls, 166 uL of the transposon mutant library (1.7 × 10^8^ CFUs) was spread on six modified KMB 1.5% agar plates in triplicate (total of 18 plates). After 24 h at 22°C the lawn was scraped off the agar surface of all six plates and collected in PBS (termed the ‘control pool’).

**Figure 1.**
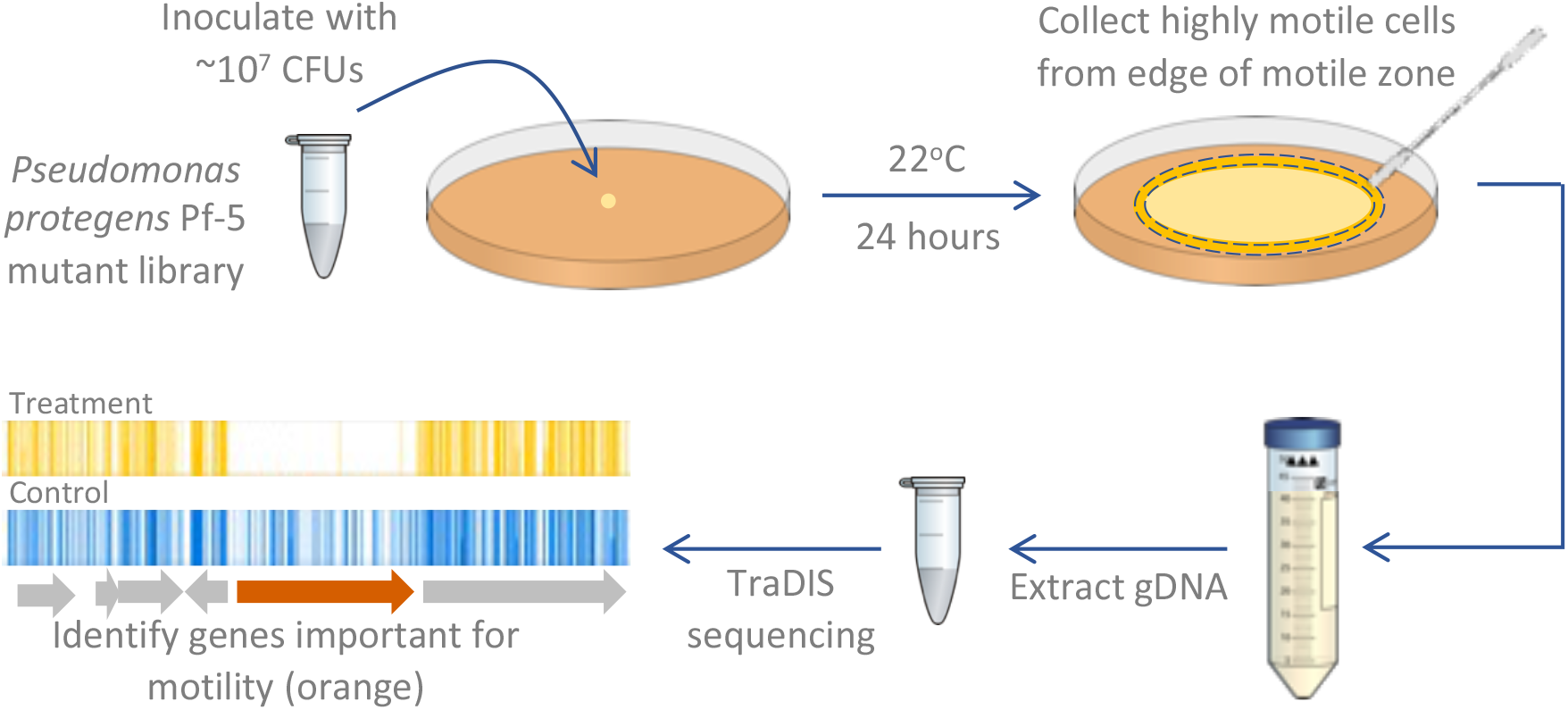
Schematic of the experimental methodology used to identify *Pseudomonas protegens* Pf-5 genes important for swimming motility. In the final panel the gene indicated in orange is important for swimming motility as the number of transposon insertion reads for this gene is significantly lower in the swimming assay compared to the control (visualized using Artemis software (36)).

### TraDIS sequencing and bioinformatic analysis

DNA was extracted from the agar/cell mixture from the output and control pools using a Wizard Genomic DNA Purification Kit (Promega). The manufacturer’s protocol was followed with two modifications: centrifugation of the precipitated DNA was increased to 18,000 x g for 5 min and to 14,000 rpm for 10 min for the ethanol-washed DNA. Agar was removed from the DNA using the Wizard SV Gel and PCR Clean-Up System (Promega). Transposon-directed insertion site sequencing (TraDIS) was performed in duplicate as previously described in Barquist et al. (33) with an Illumina MiSeq platform to obtain 52 bp single-end genomic DNA reads. Transposon insertion sites were mapped to the Pf-5 genome and analysed using the Bio-Tradis pipeline (33) as previously described (35). Briefly, this included allowing a 1bp mismatch in the transposon tag, excluding transposon insertions in the extreme 3’ end of each gene, and mapping reads with more than one mapping location to a random matching location. After matching the transposon tag, an average of 1.56 million reads per replicate were mapped to the Pf-5 genome (Table S1). A linear regression of the gene insertion indexes of the replicates was completed in R (37). Correlation coefficients between the insertion indexes for all pairs of replicates were > 0.92 (p < 0.01; Figure S1) which validates the reproducibility of our replicates and is consistent with the reproducibility of transposon insertion sequencing replicates in other studies (38). The transposon insertion sites in each of the output pools were compared with those of the control pool with the tradis_comparison.R script with default parameters. Only genes with greater than 10 reads in both replicates of either the control or output pools were included to avoid genes being falsely classified as important for fitness. Genes with a log_2_-fold change of 2 in the number of transposon insertion reads in the output pool compared to the control pool and a q-value < 0.01 were used for further analysis (39-41).

Clusters of Orthologous Groups (COG) assignments (42) for each Pf-5 gene were compiled using eggNOG-mapper (43) with 87.3% of Pf-5 coding genes assigned a COG code (35). The sum of all categories does not equal the total number of genes of interest, as some genes are assigned multiple COG codes, and some no code. Orthologs of Pf-5 genes were identified using the Pseudomonas Genome Database available at https://pseudomonas.com (44).

### Construction of gene knockout mutants

In-frame chromosomal gene deletion mutants were generated for Pf-5 genes *pstS* (PFL_6119), *phoB* (PFL_6108) and *phoR* (PFL_6109) via an overlap-extension PCR method, followed by allelic exchange with the suicide vector pEX18Tc (45), using a protocol adapted from Kidarsa et al. (46). To create the mutant allele, fragments of 500-1100 bp flanking upstream and downstream of each gene of interest were first PCR amplified using the upstream (UpF/UpR) or downstream (DnF/DnR) primer pair. All PCRs were performed using KOD Hot Start DNA polymerase (Novagen) according to (46), using the primers listed in Table S2. The upstream forward primer and downstream reverse primer each had a 5’ extension adding an *Xba*I restriction site, while the upstream reverse and downstream forward primers were designed to be in-frame with the gene of interest and had a 5’ linker of 12 bp complementary to each other to allow overlapped annealing during the secondary PCR. The amplicons of the upstream and downstream primary PCRs were gel-purified, mixed 1:1 (50 ng each) and used as the template for the secondary PCR using the UpF and DnR primers. The resultant full-length product was gel-purified, digested with *Xba*I, treated with Calf Intestinal Alkaline Phosphatase (New England Biolabs), and then cloned into the pEX18Tc vector (linearised with *Xba*I) using T4 ligase (New England Biolabs).

The recombinant vectors were transformed by electroporation into ElectroMAX DH5α-E competent *E. coli* cells (ThermoFisher Scientific) and mutant alleles verified using pEx18Tc sequencing primers (Table S3). All vectors were subsequently electrotransformed into mobilising strain *E. coli* S17-1 competent cells (47). Biparental matings were performed between the vector-containing *E. coli* S17-1 and parental Pf-5 strain for conjugative transfer of each vector into Pf-5, as described in (48), but using nutrient agar containing 1.5% (v/v) glycerol. Transconjugant Pf-5 colonies were selected on KMB agar containing 200 µg mL^-1^ tetracycline (vector conferred resistance) and 100 µg mL^-1^ streptomycin (innate resistance of Pf-5). Surviving colonies were grown without selection in LB broth for 3 h with shaking and plated on 10% sucrose LB agar to resolve merodiploids (counter-selection against *sacB*-carrying cells). Sucrose-resistant colonies were patched in parallel onto LB agar containing 10% sucrose and KMB agar with 200 µg mL^-1^ tetracycline to further confirm the absence of the pEX18Tc vector backbone. Tetracycline-sensitive colonies were screened by PCR with primers annealing to chromosomal regions external to the target gene (Table S3) to detect mutants with truncated amplicon sizes compared to a parental strain control. The deletion of each gene was confirmed by PCR and sequencing of genomic DNA from each Pf-5 mutant colony before stored in 25% glycerol at -80°C until required.

Growth curves were conducted to check for general growth defects in knockout mutants. Overnight cultures of wildtype Pf-5 and the mutant strains ∆*pstS*, ∆*phoB* and ∆*phoR* were each grown at 27°C with shaking for 16 hours in modified KMB. The cultures were sub-cultured 1:25 into fresh modified KMB and incubated with shaking until OD^600^ = 0.6.

Cultures were kept on ice while serial dilutions were performed in modified KMB to reach a final density of 1.2 × 10^4^ CFU/mL in 150 µl modified KMB in a 96-well plate (4 replicates per strain). Plates were incubated in a Pherastar plate reader at 27°C with shaking at 200 rpm and OD_600_ readings were taken at 6-minute intervals for 24 hours.

### Phenotypic assays with knockout mutants

Swimming assays were performed on a modified BM2 minimal medium (0.5% w/v casamino acids, 2 mM MgSO_4_, 10 µM FeSO_4_, 0.4% w/v glucose) supplemented with potassium phosphate buffer (0.05 or 6.6 mM; pH 7) (19) and 0.3% Bacto agar. To prepare the inoculum, cells of the Pf-5 parental strain and each mutant were scraped from a culture grown on KMB agar for 48 h, then resuspended in water and diluted to an OD_600_ of 0.2. Two microlitres of the cell suspension was stab-inoculated into the centre of swim agar plates (n = 3-4). All cultures were incubated at 22°C for 20 hr and the diameter of the motile zones were measured. We analysed the data using an unbalanced two-way ANOVA (type III) with Tukey HSD post hoc analysis using R packages car (49) and agricolae (50).

A droplet collapse assay was used to test for surfactant activity of the Pf-5 knockout mutants (51). Cultures were grown for 24 hr at 27°C on modified BM2 agar plates (as above). The cultures were scraped from the plates and suspended in water to a final density of 1 × 10^10^ CFU mL^-1^ and 10 μl droplets were spotted on parafilm in triplicate. A flat droplet indicated the cells produced the surfactant orfamide A, while domed droplets indicated the cells did not produce orfamide A (52).

Data availability. Sequence data from Transposon-directed Insertion-site Sequencing (TraDIS) are available from the European Nucleotide Archive. Sequence files for the project are available under the project accession number PRJEB56281 (https://www.ebi.ac.uk/ena/browser/view/PRJEB56281), with the control sequencing reads under sample accession numbers ERS13490282 and ERS13490283 and swimming sequencing reads under sample accession numbers ERS13490284 and ERS13490285.

## Results and Discussion

### Identifying genes contributing to Pf-5 swimming fitness

We grew a *Pseudomonas protegens* Pf-5 saturated transposon mutant library in swimming agar (0.3%) for 24 hours at 22°C and collected the cells at the edge of the motile zone (output pool). DNA from these cells was subjected to TraDIS sequencing and the read frequency for each gene was compared with that of Pf-5 cells grown on 1.5% agar (control pool). These swimming assays identified a set of 249 genes for which loss of function affected Pf-5 swimming fitness (4.40% of non-essential genes; 35; Figure 2). Loss of function of 189 of these genes were strongly detrimental for swimming fitness, based on cells with mutations in these genes being present at significantly lower levels in the output pool compared to the control (log_2_ fold change < −2). Loss of function of the remaining 60 genes was beneficial for swimming fitness, with cells carrying mutations in these genes present at significantly higher levels in the output pool compared to the control (log_2_ fold change > 2). The fold change values for the full set of Pf-5 genes are available in Dataset S1.

**Figure 2.**
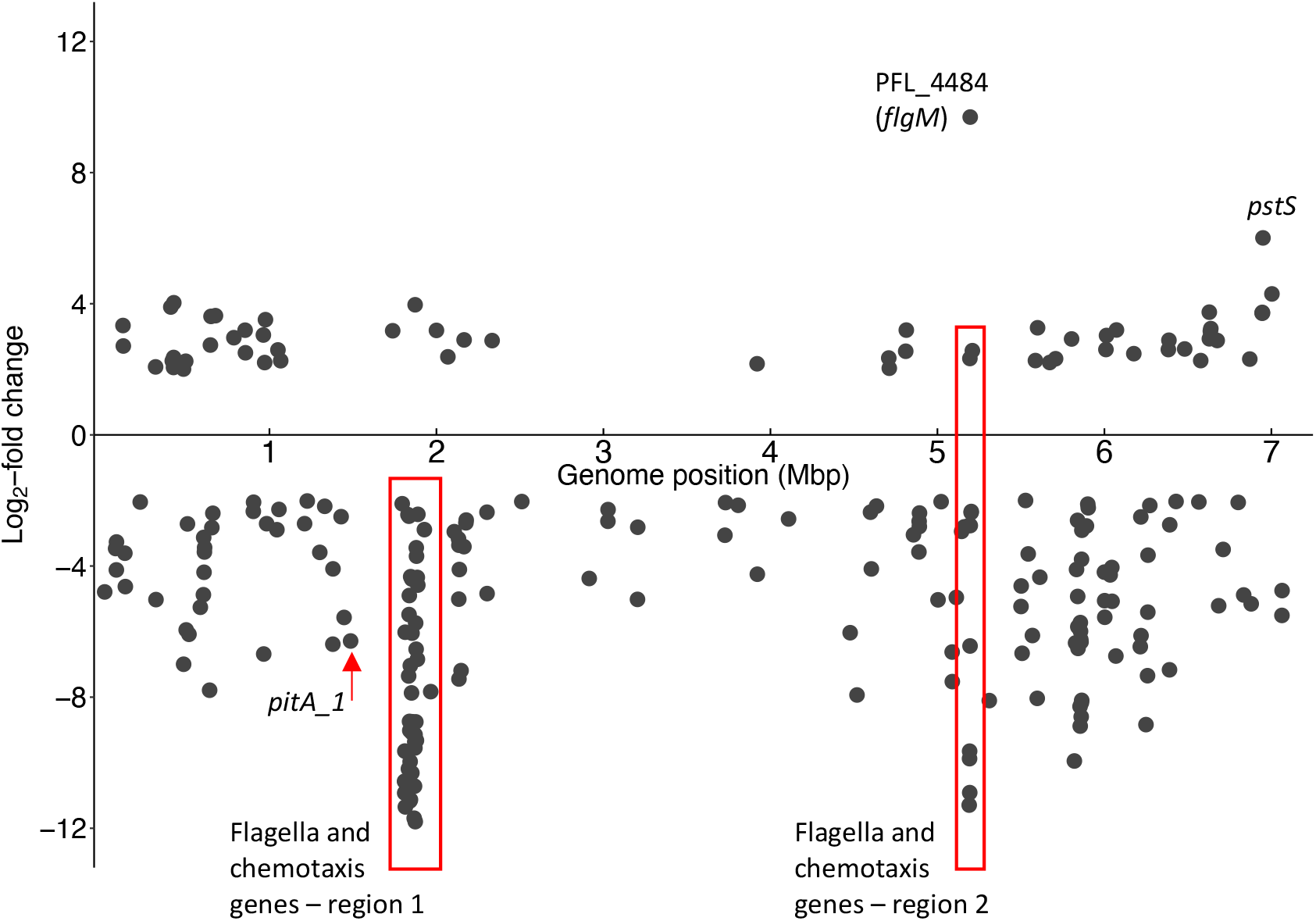
Genome position of *P. protegens* Pf-5 genes identified by TraDIS with log_2_-fold change > 2 or < -2 from swimming assays when compared with the control. Genes with a log_2_-fold change of less than -2 indicate that loss of their function was associated with reduced swimming fitness and greater than 2 indicate that loss of their function was associated with enhanced swimming fitness. Scatterplot generated using the R package ggplot2 (53).

### Functional analysis of genes contributing to swimming fitness

A functional overview of the Pf-5 genes that affected motility fitness was obtained by classifying the genes using Clusters of Orthologous Groups (COG) categories (42). As expected, genes annotated with the functional categories of cell motility (N) and signal transduction (T) made up a large proportion of the genes that detrimentally affected swimming fitness when their function was lost (30.7%; Figure 3). The functional loss of a large proportion of the genes in the following categories also detrimentally affected swimming motility: cell wall/membrane/envelope biogenesis (M); post-translational modification, protein turnover and chaperones (O); energy production and conversion (C); replication, recombination and repair (L); and unknown function (S; Figure 3). From the set of genes that enhanced swimming fitness when their function was lost, the highest proportion are involved in signal transduction (T), followed by translation, ribosomal structure and biogenesis (J), inorganic ion transport and metabolism (P), and function unknown (S; Figure 3). Just over 20% of this gene set have no COG code assigned, including pseudogenes, tRNAs, rRNAs and conserved hypothetical proteins.

**Figure 3.**
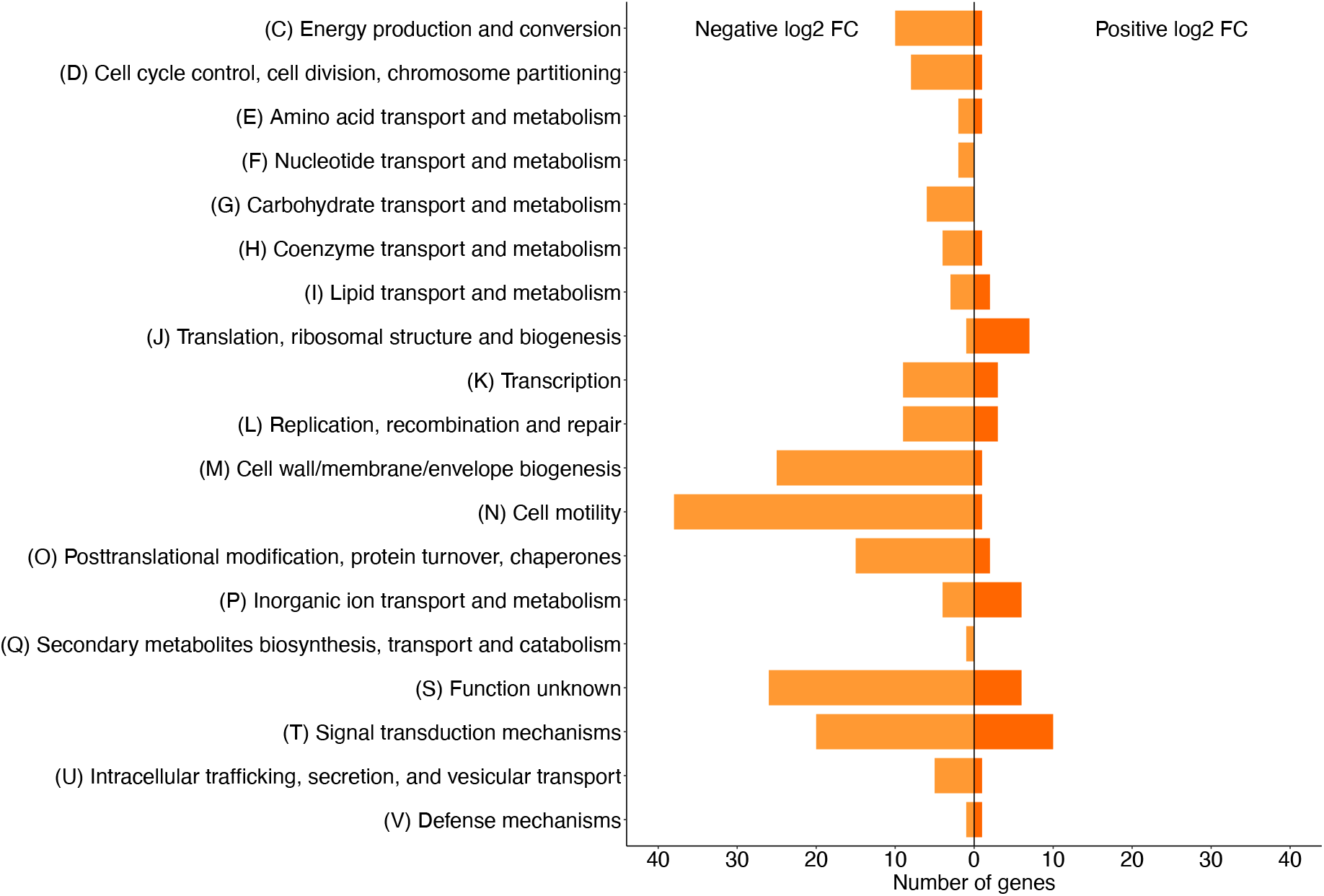
Functional classification of genes that affect swimming fitness when their function is lost (log_2_ fold change > 2 or < -2). Data visualized using the R package ggplot2 (53).

### Swimming motility requires a large number of flagellar and chemotaxis genes

Of 48 genes related to flagella structure, biosynthesis, assembly, and regulation, which are mainly located in two regions of the Pf-5 genome, our TraDIS analysis identified 41 as significantly important for swimming motility (Figure 4). In addition, loss of 18 chemotaxis genes affected Pf-5 swimming fitness (Figure 4). The direction of the fold changes for the flagella genes and most of the chemotaxis genes are consistent with the known roles of orthologous proteins in *P. aeruginosa*, but there are a few chemotaxis genes that affected Pf-5 swimming fitness in an unexpected way (*pctA_2*, PFL_0778, *cheZ*, PFL_4482, PFL_5046).

**Figure 4.**
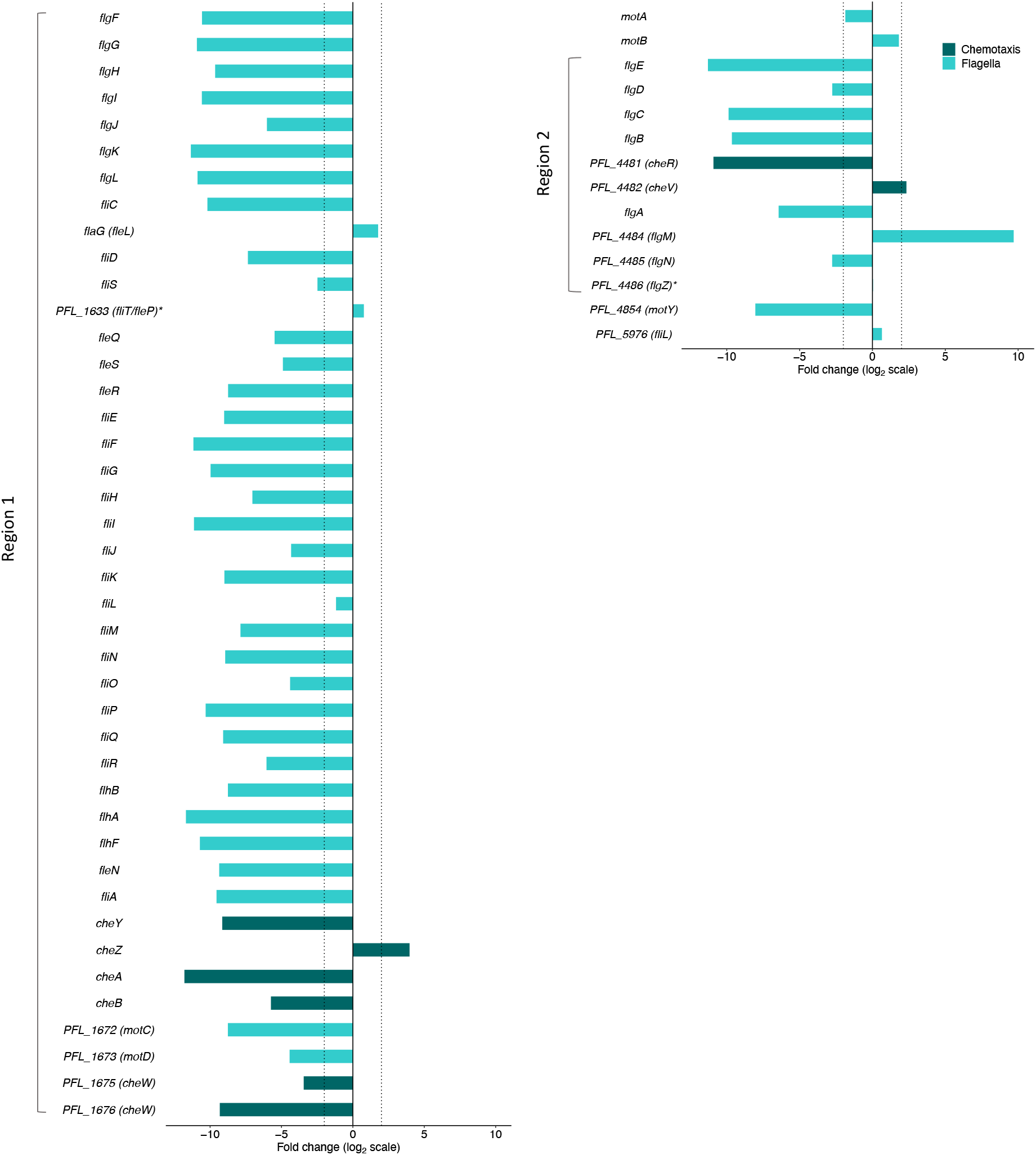
Log^2^ fold change of *P. protegens* Pf-5 flagella and chemotaxis cluster I and V genes during swimming (not including genes that encode methyl-accepting chemotaxis proteins). Genes are clustered in two main regions of the genome, indicated as regions 1 and 2. Dotted vertical lines are positioned at log_2_ fold changes of -2 and 2. Gene names from *P. aeruginosa* orthologs in brackets. * genes with non-significant log_2_ fold change. Data visualized using the R package ggplot2 (53).

The chemosensory genes of Pf-5 are organized in a similar way to the five *P. aeruginosa* PAO1 gene clusters that encode chemosensory signaling proteins, except there are no cluster II orthologs, only some orthologs of cluster IV genes are present and *chpA* is truncated (54, 55). Loss of most of the Pf-5 genes orthologous to those in *P. aeruginosa* clusters I and V (PFL_1668-PFL_1677 and PFL_4481-PFL_4482, respectively) detrimentally affected swimming (Figure 4). In *P. aeruginosa* the Che pathway, comprised of cluster I and V genes, is essential for chemotaxis, which is consistent with the results of this study (56). In contrast, loss of function of *cheZ* (cluster I) was beneficial for Pf-5 swimming. Whereas, in *E. coli*, loss of *cheZ* function results in a state of tumbling and random movement (57, 58). Loss of function of PFL_4482 (ortholog of *cheV* in cluster V) was also beneficial for swimming. The exact role of CheV in bacterial cells is not well understood, but it possibly acts as a phosphate sink in enterobacteria by competing with CheY for phosphorylation by CheA to offset chemoreceptor overstimulation (59).

In Pf-5 there are 42 genes that encode known chemosensory receptors, comprising the named genes *pilJ, pctC, pctA_1, bdlA, aer_1, aer_2, aer_3* and *pctA_2*, and a further 34 which have no specific assigned function and are annotated only as methyl-accepting chemotaxis proteins (MCPs). MCPs are critical components of chemosensory signaling systems, but the majority are not located in a cluster or pathway (54). Three of the Pf-5 genes encoding MCPs influenced swimming fitness when disrupted: loss of function of PFL_0778 and *pctA_2* was detrimental for swimming fitness, whereas loss of PFL_5046 function was beneficial for swimming fitness. The chemoattractants of these three genes remain unknown. Loss of function of *aer_2*, which encodes an aerotaxis receptor, detrimentally affected Pf-5 swimming when disrupted. This gene is homologous to the *aer* gene in *P. aeruginosa* PAO1 that encodes the most prevalent aerotaxis receptor in pseudomonads (60). Consistent with our results, loss of the gene encoding the main aerotaxis receptor in the biocontrol bacteria *P. chlororaphis* PCL1606 detrimentally affected swimming motility (61).

### Defects in cell envelope and cell division genes are detrimental for swimming motility

There are 43 cell envelope and cell division genes that detrimentally affected Pf-5 swimming fitness when their function was lost (Figure 5a and 5b). Mutations in these genes are likely to affect cell shape or cell surface composition (62-64). Swimming motility is based on cells sensing environmental cues and switching between ‘runs’ and tumbling movement to move up or down a chemical gradient (6). Changes in cell morphology have the potential to impact on the drag and other forces experienced by the cell and therefore their swimming fitness (65). For example, in *E. coli* AW405 changes in cell length are linked to changes in swimming patterns, and *E. coli* KR0401 *rodZ* mutants defective in peptidoglycan synthesis are spherical (instead of the normal rod shape) and non-motile (66-68).

**Figure 5.**
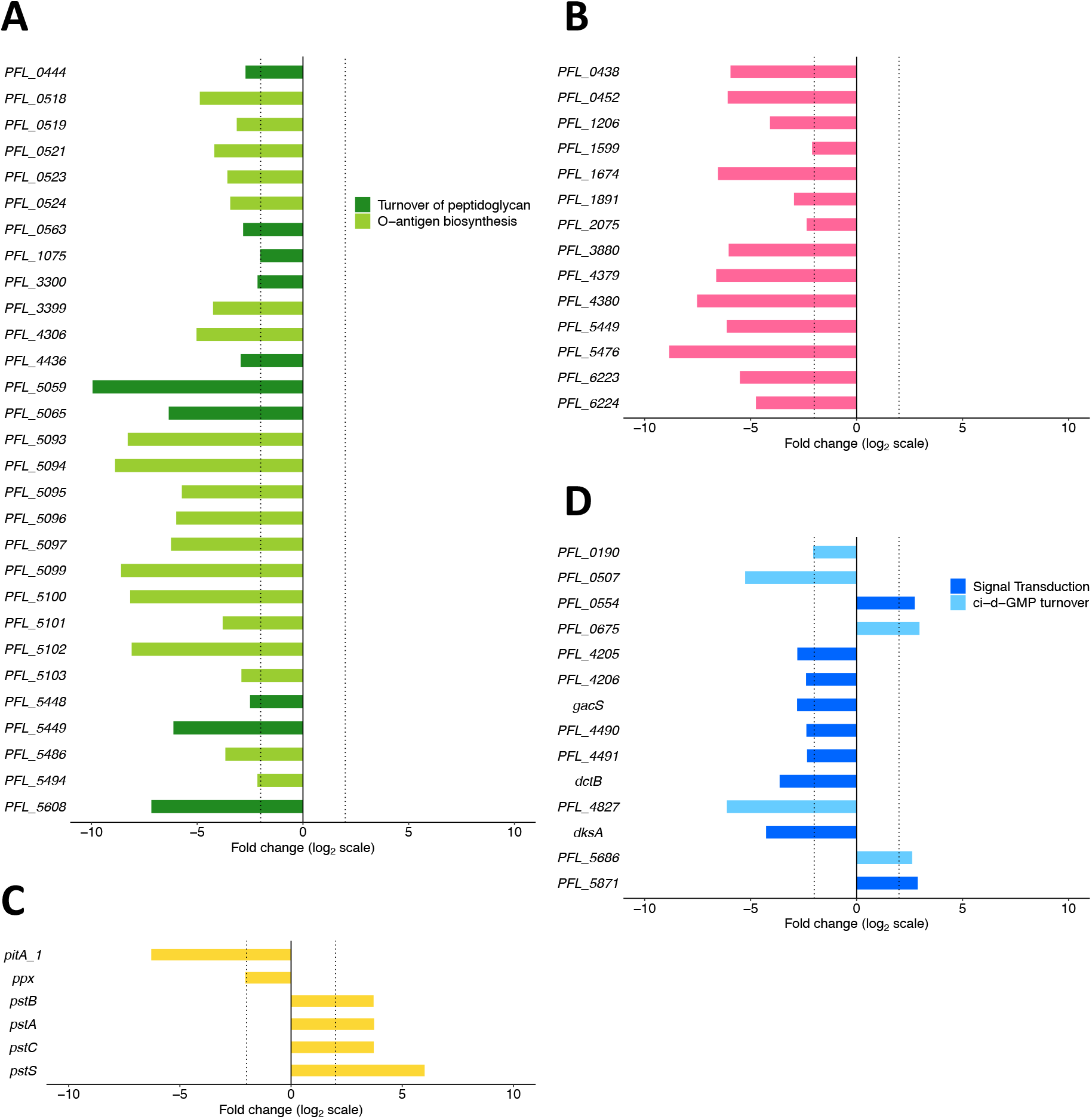
Log_2_ fold change of *P. protegens* Pf-5 genes related to (A) cell envelope, (B) cell division, (C) phosphate turnover and (D) signal transduction during swimming. Dotted vertical lines are positioned at log_2_ fold changes of -2 and 2. All genes have significant log_2_ fold change. Data visualized using the R package ggplot2 (53).

Nineteen of the 43 cell envelope genes have functions in lipopolysaccharide (LPS) biosynthesis, including genes associated with LPS core biosynthesis (PFL_0518, PFL_0519, PFL_0521, *wapB, galU, wzt*) as well as those related to O-antigen biosynthesis (*wbpL, rfbG, rfbH*). Mutations causing defects in LPS are likely to affect cell surface structures and have previously been shown to affect motility in multiple bacterial species, including *P. aeruginosa* PAO1 and *P. syringae* pathovar *glycinea* (69, 70). Impacts of LPS defects on swimming motility have been shown to be due to changes in flagella and/or changes in cell adhesive properties. In *E. coli* and *S. enterica* serovar Typhimurium the expression of flagellar genes is impaired by cell envelope stress, specifically the truncation of LPS (71). In *P. aeruginosa* PAO1 mutants with non-wildtype LPS have reduced swimming motility due to changes in cell-cell and cell-substrate adhesion forces (72).

Ten of the 43 cell envelope-related genes are involved in peptidoglycan turnover (Figure 5a). This includes genes from each stage of peptidoglycan turnover, from synthesis of muropeptides (*ddlB* and *murF*), to incorporating new muropeptides into existing peptidoglycan (*mrcA*, PFL_3300, PFL_5449, and PFL_5608) and modifying/degrading the peptidoglycan (*dacA, mltF*, PFL_0563 and PFL_4436). The loss of function of four of these genes had very strong negative effects on swimming fitness (log_2_ fold change < -4), indicating that the correct assembly of the precursor molecule D-alanyl-D-alanine by DdlB and MurF, and insertion of new muropeptides into the cell wall are critical for swimming fitness. In *P. aeruginosa* PAO1 and *E. coli*, the proteins encoded by *dacA* and PFL_5449 (RlpA family lipoprotein) have been shown to be important for maintaining cell shape (73, 74). It is therefore likely that the effect of the loss of peptidoglycan turnover-related genes on Pf-5 swimming fitness is through alterations to typical cell shape.

Thirteen cell division-related genes detrimentally affected swimming fitness, with 10 having a strong negative effect (log_2_ fold change < -4; Figure 5b). These genes include those involved in localizing the cell division machinery to the center of the cell (*minC* and *minD*) and breaking down peptidoglycan for septal splitting (*nlpD*) along with genes involved in segregation of origin of replication domains (*smc, scpA*), chromosome or plasmid molecules (*parA, parB*), and the chromosome terminus (*ftsK*). Loss of function of cell division genes likely results in abnormal cell morphology. For example, *minC* and *minD* mutants in the plant pathogen *Xanthomonas oryzae* pv. *oryzae* form short filaments and minicells and a *minC* knockout was not able to swim (75). *P. aeruginosa* PAO1161 *parA* and *parB* mutants also have swimming defects (76, 77).

### Swimming motility is affected by defects in signal transduction genes

Loss of function of 14 signal transduction genes affected swimming fitness, including the sensor kinase *gacS* which is part of the global regulatory GacSA signal transduction system in pseudomonads (Figure 5d; 29). In this study loss of *gacS* function was detrimental for Pf-5 swimming motility. In other pseudomonads the effect of *gacS* mutation on swimming motility has been shown to be varied. For example, *gacS* mutants of *P. aeruginosa* PA14, *P. chlororaphis* O6 and *P. fluorescens* F113 showed enhanced swimming compared to wildtype (78-81). In contrast, a *gacS* mutant of *P. fluorescens* NBC275 had decreased motility compared with the wildtype strain (21).

The loss of function of five genes involved in c-di-GMP turnover affected Pf-5 swimming fitness. Turnover of the second messenger c-di-GMP controls a range of cellular processes, including motility, and a low concentration of intracellular c-di-GMP is associated with motile cells (82). The loss of PFL_4827, an ortholog of *bifA* that encodes a phosphodiesterase in *P. aeruginosa* PAO1, was detrimental for Pf-5 swimming.

Phosphodiesterases are part of the c-di-GMP regulation system in pseudomonads and BifA has been shown to degrade c-di-GMP in *P. aeruginosa* PA14 (83). In *P. fluorescens* F113 and *P. putida* KT2440, *bifA* knockouts had reduced c-di-GMP breakdown, causing higher c-di-GMP levels and a reduction in swimming motility (84, 85). The loss of function of PFL_0190 and PFL_0507 also had detrimental effects on Pf-5 swimming motility. These genes are annotated as diguanylate cyclase proteins, but in *P. aeruginosa* PAO1 the ortholog of PFL_0507, *dipA*, functions as a phosphodiesterase and knocking it out resulted in reduced motility (86). Enhanced Pf-5 swimming fitness is seen with the loss of function of PFL_0675 and PFL_5686 which encode diguanylate cyclase proteins that synthesize c-di-GMP. In a separate Pf-5 study, a PFL_5686 mutant showed wildtype swimming motility (87). In *P. aeruginosa* PAO1 and *P. fluorescens* Pf0-1 *gcbA* knockouts had increased swimming motility compared to the wildtype strain (88, 89).

### Swimming motility also requires conserved genes of unknown function

Loss of function of 24 genes encoding conserved hypothetical proteins affected Pf-5 swimming fitness (Dataset S1), with six of these genes enhancing swimming fitness, and the other 18 genes negatively affecting swimming fitness. Loss of function of 13 genes encoding conserved hypothetical proteins had a very strong effect on swimming fitness (log_2_ fold change > 4 or < -4) so characterization of these genes is of particular interest and may provide further insights into swimming motility in this plant-associated bacteria.

Genomic context and ortholog function analysis suggested possible functions for five hypothetical genes. PFL_1677 is in an operon with two CheW-like genes (PFL_1675–1677), suggesting a possible role in chemotaxis, and loss of function of all three genes in the operon detrimentally affect swimming fitness. A further two are likely cell envelope genes (PFL_5098, PFL_5485). PFL_5098 is part of a cluster of 11 O-antigen biosynthesis genes (PFL_5093–5103) that all have detrimental effects on swimming fitness when lost. PFL_5485 is in an operon (PFL_5485–5488) which also contains genes encoding a GtrA family protein and a glycosytransferase. In *P. donghuensis* HYS GtrA is involved in the glucosylation of lipopolysaccharide O-antigen (90). There is also evidence that PFL_1916 may be associated with energy transduction as it is in a two-gene operon with *ccoG_1* which encodes a cytochrome oxidase assembly factor.

Conserved hypothetical proteins comprise all (or the majority of) genes of two operons (PFL_2736–2739 and PFL_5239–5241) where loss of each of the members were detrimental for Pf-5 swimming fitness, but genomic context and ortholog function analysis have not shed any light on their potential functions. Now that we have connected these genes with swimming motility, this may assist in future functional characterization.

### A role for phosphate transport in Pf-5 swimming fitness

A range of genes involved in phosphate utilization showed differential effects on swimming fitness. Loss of function of *pitA_1*, encoding a low-affinity inorganic phosphate transporter, and *ppx*, encoding exopolyphosphatase, were detrimental for swimming fitness (Figure 5c). In *P. aeruginosa* PAO1, *ppx* encodes an exopolyphosphatase which cleaves P_i_ residues from accumulated polyP chains. In *P. aeruginosa* PAO1 a *ppx* mutant had reduced swimming motility, consistent with the Pf-5 TraDIS data (91).

Interestingly, swimming fitness increased when the genes encoding the Pst high-affinity inorganic phosphate transporter (*pstBACS*) were lost, with *pstS* showing the second greatest fold increase across the genome (Figure 5c). In *P. aeruginosa* this transporter is active in low-phosphate conditions and is part of a wider system of sensing and responding to environmental inorganic phosphate conducted by the Pho regulon (92), but has not previously been connected to swimming. We made gene knockouts in the transporter gene *pstS*, and the regulatory genes *phoB* and *phoR* (not significantly affected in the TraDIS swimming dataset) to investigate the role of the Pho regulon in swimming. We conducted swimming assays with the three mutants and the parental strain under phosphate-replete (6.6 mM) and phosphate-deplete conditions (0.05 mM). Under both phosphate levels !ι*pstS* had a significantly larger swimming zone diameter than the parental strain whereas !ι*phoB* and !ι*phoR* showed the same swimming phenotype as the parental strain (F(9,39) = 2.32, p = 0.03; Figure 6 and S3) consistent with our TraDIS results. To confirm increased swimming motility was not due to overall growth improvements in these mutants, we conducted growth assays. Mutants of *phoB* and *phoR* showed identical growth curves to the parental strain, but the *pstS* mutant had slower growth in liquid culture compared to the parental strain, indicating swimming improvements were not due to an enhanced growth rate (Figure S4). A droplet collapse assay was conducted and showed that production of the biosurfactant orfamide A was identical between the mutants and parental strain (52). The biosurfactant orfamide A has previously been shown to be involved in swarming motility in Pf-5 (93). Overall, both the gene knockouts and TraDIS data are consistent with PstS playing an important role in swimming motility in Pf-5, whereas the regulators PhoB and PhoR do not affect swimming motility. Expression of *pstS* is highly induced under phosphate deplete conditions in *P. aeruginosa* PAO1 (94) so it is interesting that in Pf-5 PstS appears similarly important for swimming motility in both phosphate replete and deplete conditions. In *P. aeruginosa* there is a link between phosphate, the Pho regulon and swarming motility (94, 95), but there are no reports of links with swimming.

**Figure 6.**
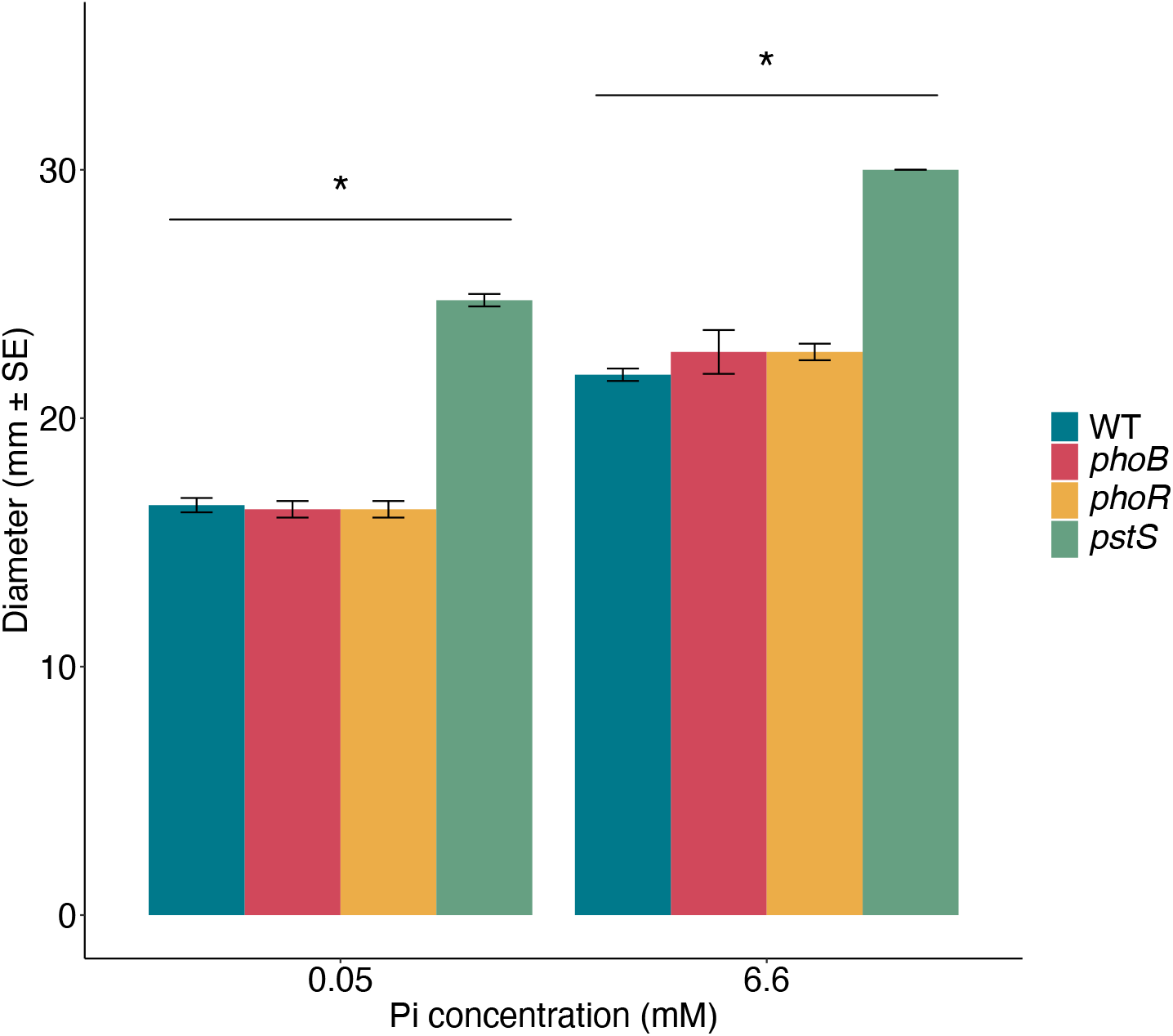
Swimming assays conducted with *P. protegens* Pf-5 single knockout mutants under two phosphate concentrations. The mean diameter (mm ± SE) of the motile zones measured after 20 hours growth at 22°C on BM2 0.3% agar plates (*phoB* and *phoR* n = 3; *pstS* and the Pf-5 parental strain n = 4). An unbalanced two-way ANOVA with Tukey HSD post hoc analysis was conducted in R (37). * indicates significant difference at α = 0.05. Figure generated using the R packages ggplot2 (53) and ggsignif (96).

## Conclusions

In this study TraDIS successfully identified 249 genes involved in swimming motility of *P. protegens* Pf-5. This method identified both genes that are known to contribute to swimming motility as well as genes not previously associated with this phenotype. Successful swimming motility required the function of chemotaxis genes, genes involved in flagella structure, assembly, and regulation, and signal transduction genes involved in c-di-GMP turnover. We also identified additional genes for which the loss of function negatively affected swimming motility, including cell division, peptidoglycan turnover and lipopolysaccharide biosynthesis genes, with effects on swimming likely due to aberrations in cell morphology. Of particular interest is our finding that phosphate transport plays a role in swimming motility. Both TraDIS and specific gene knockout results link phosphate transport by the Pit and Pst transporters with swimming motility in Pf-5, a connection not previously reported. Phosphate is an essential nutrient for cell function, and as phosphate is often limited in agricultural soils the link between phosphate transport and motility in biocontrol bacteria could be crucial for enabling biocontrol activity and preventing crop disease.

Overall, this study expands our knowledge of pseudomonad motility and highlights the utility of a TraDIS-based approach for systematically analyzing thousands of genes and identifying the extended suite of genes affecting the fitness of the plant growth promoting rhizobacteria Pf-5 during swimming. Understanding the genetic basis of swimming motility enhances our knowledge of key processes in biocontrol bacteria that are needed to ensure their competitive success. This knowledge will help address the issue of inconsistency in biocontrol bacteria effectiveness and reliability in the field and contribute to developing strategies to increase use of biocontrol bacteria in agricultural settings to prevent crop losses.

## Acknowledgements

The TraDIS sequencing for this study was conducted at The Ramaciotti Centre for Genomics at the University of New South Wales. *Pseudomonas protegens* Pf-5 was obtained from Professor Joyce Loper (Oregon State University). We thank Dr Liam Elbourne (Macquarie University) for setting up the local installation of the Bio-Tradis bioinformatics pipeline, Dr Amy Cain (Macquarie University) for advice on TraDIS experimental design, Dr Francesca Short (Monash University) for advice on TraDIS data analysis, Dr Qing Yan (Oregon State University) for technical advice on the construction of knockout mutants and Professor Joyce Loper (Oregon State University) and Dr Virginia Stockwell (USDA) for valuable discussions and advice. This work was supported by an Australian Research Council Discovery Grant (DP160103746). BF is the recipient of an Australian Government Research Training Pathway Scholarship and KAH is supported by an Australian Research Council Future Fellowship (FT180100123).

