## Supplementary tables and figures for "Identifying the suite of genes central to swimming in the biocontrol bacteria *Pseudomonas protegens* Pf-5"

**This PDF file includes:**

Table S1

Table S2

Table S3

Figure S1

Figure S2

Figure S3

Figure S4

**Other supplementary materials for this manuscript include:**

Dataset S1

Table S1. *Pseudomonas protegens* Pf-5 transposon mutant library metrics from Bio-Tradis pipeline analysis.

| **Treatment** | **Replicate** | **Total no. of reads** | **No. (%) of reads  with matching  transposon tag** | **No. (%) of aligned reads** | **No. of unique insertion sites** | **Average distance between unique insertion sites (bp)** |
| --- | --- | --- | --- | --- | --- | --- |
| Swimming | 1 | 1,363,792 | 1,359,791 (99.7%) | 1,333,222 (98.0%) | 306,381 | 23.1 |
| Swimming | 2 | 1,758,995 | 1,753,807 (99.7%) | 1,718,672 (98.0%) | 322,915 | 21.9 |
| Control | 1 | 3,951,744 | 3,922,215 (99.3%) | 2,187,032 (55.8%) | 352,775 | 20.1 |
| Control | 2 | 2,790,059 | 2,770,318 (99.3%) | 1,941,521 (70.1%) | 345,809 | 20.5 |

**Supplementary tables**

Table S2. **Primers for allelic exchange mutagenesis of genes. The random six nucleotides are italicised, restriction enzyme recognition sites are indicated in bold, and the complementary regions between the UpR/DnF primers are underlined.**

| **Genes** | **Amplifi-cation regions** | **Primer names** | **Primer sequences (5’-3’)** | **Amino acids remaining (incl start & stop codon)** |
| --- | --- | --- | --- | --- |
| PFL_6108 | 5’ flanking | 6108-UpF-XbaI | *CGTGCT***TCTAGA**GGAGGTCAGTCGGATTTCGC | 10 |
|  |  | 6108-UpR | GGCCTTGGTGGAGCTTCTGCCAACCATGCTTG |  |
|  | 3’ flanking | 6108-DnF | GTTGGCAGAAGCTCCACCAAGGCCTGACGG |  |
|  |  | 6108-DnR-XbaI | *GTGAGG***TCTAGA**CGTAGCTGTCCTGCTCGAAG |  |
| PFL_6109 | 5’ flanking | 6109-UpF-XbaI | GTGAGG**TCTAGA**ATCATCATGCTCACCGCCAA | 18 |
|  |  | 6109-UpR | GGTCAGGCTCTGAATCAGGGTGCCATGCCAGT |  |
|  | 3’ flanking | 6109-DnF | GGCACCCTGATTCAGAGCCTGACCGACCAG |  |
|  |  | 6109-DnR-XbaI | *ATGACG***TCTAGA**GACCACGATGTGCAGGTAGG |  |
| PFL_6119 | 5’ flanking | 6119-UpF-XbaI | *ATGACG***TCTAGA**GCTCTTCCAGGGAGAACATC | 9 |
|  |  | 6119-UpR | GGCATGAAACTGCTGAAAGAAGGCGCATAAGC |  |
|  | 3’ flanking | 6119-DnF | GCCTTCTTTCAGCAGTTTCATGCCTTACTCCT |  |
|  |  | 6119-DnR-XbaI | *GTGAGG***TCTAGA**GTTCGGCAGCGTCTTGTTAC |  |

Table S3. **Primers for verification of gene deletions via PCR and sequencing. These primers anneal external to the gene deletion region of the genome, or within the vector backbone external to the cloned insert region.**

| **Genes** | **Functions** | **Primer names** | **Primer sequences (5’-3’)** |
| --- | --- | --- | --- |
| PFL_6108 | Verification of Δ*phoB* mutant | 6108ExtF | CGTCAAGCAAGGCCCTATGG |
|  |  | 6108ExtR | TTGCGCATCTGTTCCAGCT |
| PFL_6109 | Verification of Δ*phoR* mutant | 6109ExtF | GACCTGATCCTGCTGGACTG |
|  |  | 6109ExtR | GGTAGATGGCCGGGTACATC |
| PFL_6119 | Verification of Δ*pstS* mutant | 6119ExtF | GGATCGGCAGGTCGACTTTG |
|  |  | 6119ExtR | ACAGCCTGGCCATTATCTTCC |
| pEx18Tc vector | Verification of mutant allele constructs | pEX18Tc_F | CCTCTTCGCTATTACGCCAG |
|  |  | pEX18Tc_R | GTTGTGTGGAATTGTGAGCG |

**Supplementary figures**

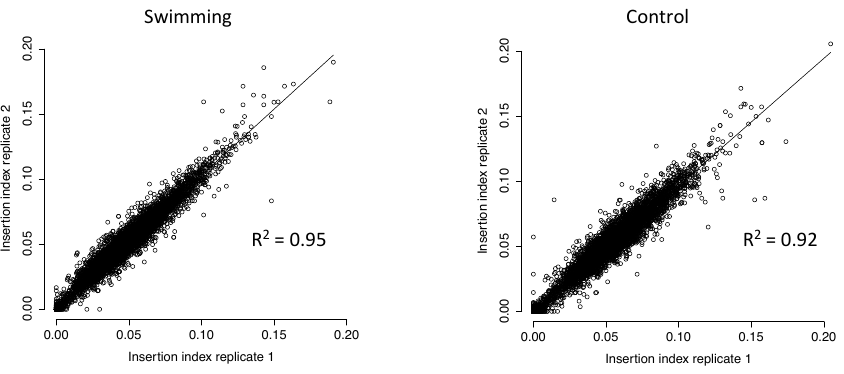

Figure S1. Correlation of gene insertion indexes for the two replicates of the *Pseudomonas protegens* Pf-5 mutant library in swimming conditions and the control. Insertion index is calculated as the number of transposon insertion sites in a gene divided by the gene length. Figure generated using R (1).

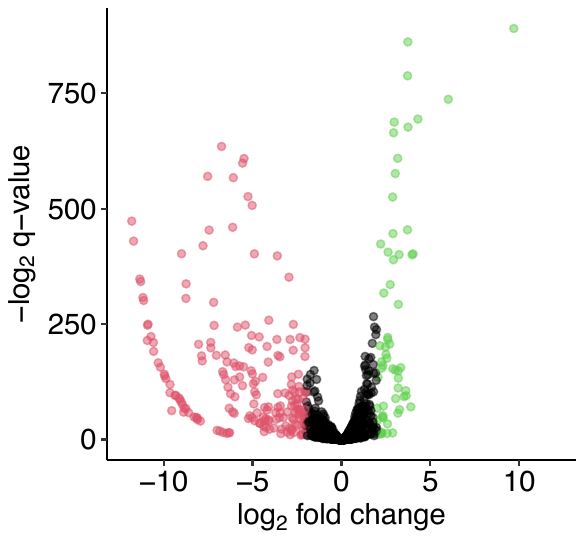

Figure S2. Volcano plot showing log_2_-fold change of all *Pseudomonas protegens* Pf-5 genes from swimming assay when compared with the control. Points in red have a significant fold change < -2 (p < 0.01); loss of these genes is detrimental for swimming fitness. Points in green have a significant fold change > 2 (p < 0.01); loss of these genes is beneficial for swimming fitness. Figure generated using the R package ggplot2 (2).

| Phosphate concentration | Parental  strain | ∆*phoB* | ∆*phoR* | ∆*pstS* |
| --- | --- | --- | --- | --- |
| 6.6 mM | 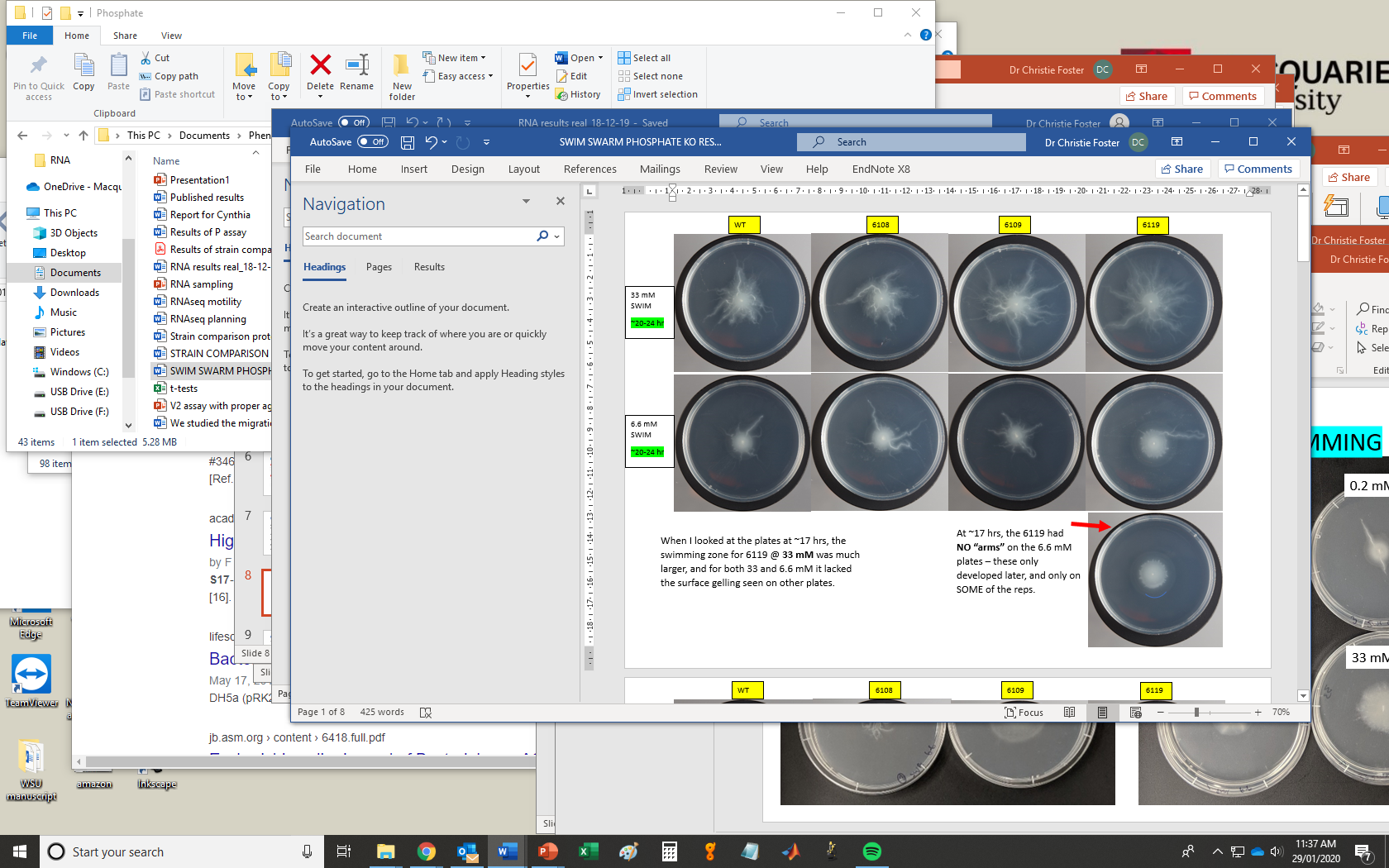 | | | |
| 0.05 mM | 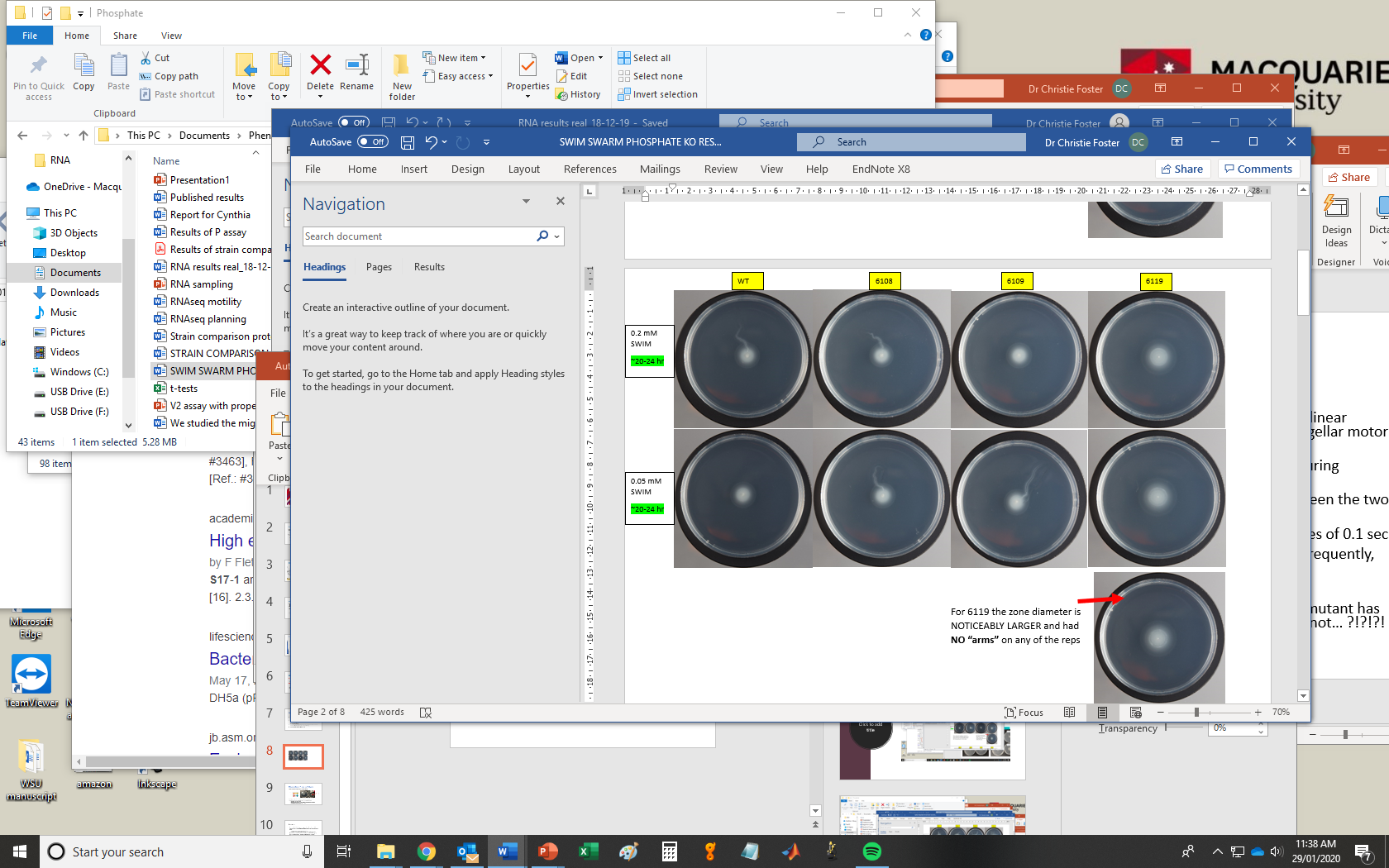 | | | |

Figure S3. Representative swimming assay plates for *Pseudomonas protegens* Pf-5 parental strain and *pstS*, *phoB* and *phoR* single knockout mutants after 24 hours growth at 22^o^C on BM2 0.3% agar plates with 0.05 or 6.6 mM phosphate.

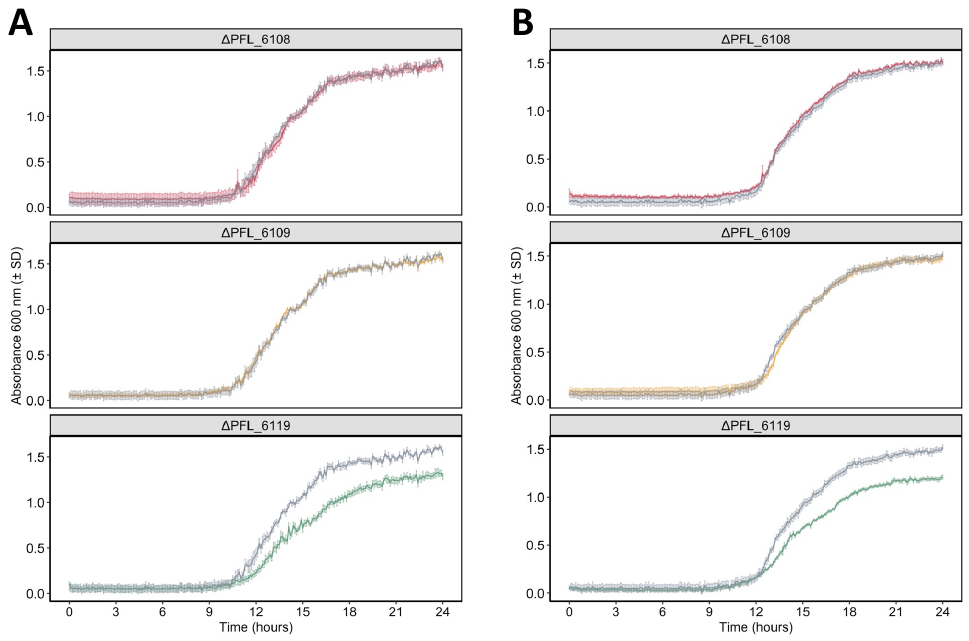

Figure S4. Growth of *Pseudomonas protegens* Pf-5 single knockout mutants compared to wild-type (grey) on (A) BM2 broth + 6.6 mM phosphate and (B) BM2 broth + 0.05 mM phosphate. Absorbance was measured at 600 nm, four replicates per strain were used and the mean absorbance (± SD) is shown. Figure generated using the R package ggplot2 (2).

**Legend for Dataset S1**

Dataset S1. Fold changes of all *Pseudomonas protegens* Pf-5 genes from comparing the output pool (motile cells) to the control pool. Metadata located in separate tab.
